# Respiratory Syncytial Virus in Infants and Children with Prader-Willi syndrome

**DOI:** 10.1101/2020.06.23.166660

**Authors:** JL Miller, E Thornton

## Abstract

**BACKGROUND:** Prader-Willi syndrome (PWS) is a complex disorder affecting approximately 1/15,000-1/30,000 people. Infants with PWS are at risk for serious complications with Respiratory Syncytial Virus (RSV) due to low muscle tone and a weakened pulmonary system.

**OBJECTIVES:** Understanding RSV incidence, hospitalization rates, lingering effects, and morbidity in children with PWS may help with planning health care, insurance and vaccine recommendations in children with PWS.

**METHODS:** Links to volunteer surveys were provided via direct email and social media to families throughout the United States with children having PWS. The contact distribution lists were provided by PWSA(USA) and the Foundation for Prader-Willi Research.

**RESULTS:** A total of 220 surveys were completed by the parents/caregivers of children with PWS. Of those respondents, 60 (27.27%) had contracted the RSV virus during early childhood. Of those with RSV, 44 children required hospitalization, with 16 reporting multiple hospitalizations, some for several weeks. Of those with the virus, 22 required PICU admission, 10 required intubation, 20 needed CPAP, and 46 children needed supplemental oxygen during the infection. Of those who had contracted RSV, 42% were over age 1 year at the time of infection, and 17 children developed chronic lung issues after the RSV infection. The case lethality was 1.37%.

Only 38% had received the RSV Synagis shot, and 19% received more than one season of the vaccination. Prematurity prevalence was only 28%, but 99% reported that their child had significant hypotonia.

Approximately 30% of parents sited lack of insurance authorization or failure of the physician to recommend the treatment.

**CONCLUSIONS:** The risk of contracting RSV for young children with PWS is high. The implications of contracting RSV include death or lung damage, along with high medical expenditures, which could be ameliorated with routine administration of the Synagis vaccine.

## INTRODUCTION

Prader-Willi syndrome (PWS) is a complex neurodevelopmental disorder that conveys hypotonia, risk of obstructive and central sleep apnea, multiple endocrinopathies, and eating issues that change between infancy and adolescence/adulthood. PWS is considered a rare disease and affects approximately 1:15,000 – 1:30,000 births [1,2]. With improved physician awareness of the syndrome as a cause of hypotonia, and improved genetic testing techniques, the diagnosis is often made within the first few months of life [1]. The biggest advantage of early diagnosis is the ability for parents to be proactive about treatments their child may need and risks they will face during life [3]. It is well recognized that early diagnosis results in access to therapies (physical, occupational, feeding) and early referral to endocrinologists for treatment with growth hormone therapy [3,4]. However, the importance of recognizing the risks associated with the syndrome should not be underestimated.

Bronchiolitis is the most common respiratory infection in the first 2 years of life, and respiratory syncytial virus (RSV) is the most frequently responsible virus [5]. In “high-risk groups,” RSV can cause severe respiratory impairment, leading to hospitalization, intensive care, and even death. Respiratory infections account of more than 50% of deaths in individuals with PWS [6]. Respiratory infections were the primary cause of death for children with PWS <1 year of age and account for 50% of deaths for children less than 2 years of age [6]. Multiple studies have confirmed that respiratory failure/infections are a common cause of mortality in young children with PWS [7,8].

Specific high-risk groups, culled out by the American Academy of Pediatrics include children with congenital heart disease, infants with neuromuscular impairment, cystic fibrosis, Down Syndrome, immunodeficiency syndromes and extreme prematurity [5]. It is documented that children who fall into these high-risk groups have a 3- to 10-fold increase in the rate of hospitalization due to RSV infection [5]. Children in these groups are at high risk for morbidity and mortality from this extremely common respiratory infection [9].

Individuals with PWS should qualify for the Synagis vaccine under the neuromuscular disease category, given the almost 100% prevalence of severe hypotonia [1]. However, in clinical practice, we observed a high incidence of hospitalizations for children with PWS who contracted RSV. Knowing that recommendations from the American Academy of Pediatrics should support all infants and young children receiving the RSV vaccine, we were surprised with the frequency of need for hospitalization and intensive care treatment in this population. This led us to investigate how many children with PWS actually receive the vaccine in infancy, as well as the relationship between not receiving the vaccine and morbidity/mortality in this population.

## METHODS

Links to volunteer surveys were provided via direct email and social media to families throughout the United States with children with PWS. The contact distribution lists were provided by PWSA(USA) and the Foundation for Prader-Willi Research. Informed consent was obtained by the PWSA (USA) and/or Foundation for Prader-Willi Research.

All survey reports were anonymous. Parents were asked a series of 24 questions on the survey regarding whether or not their child had had RSV, need for hospitalization, intubation during infection, whether or not they had received the Synagis vaccine, and if no vaccine, why not, as well as whether or not their child died from RSV. Genetic subtype, medical history of lung issues, prematurity, and hypotonia were queried, in addition to the age at time of RSV infection.

## RESULTS

A total of 220 surveys were completed by the parents/caregivers of children with PWS (50% male/50% female). Of those respondents, 60 of 220 children (27.27%) of the children with PWS had been confirmed to have the RSV virus (Figure 1). Forty-four of those children with RSV required hospitalization (73%), with 16 reporting multiple hospitalizations, some for several weeks. Of those who were hospitalized, 50% required admission to the intensive care unit and 10 of those children required mechanical ventilation.

**Figure 1:**
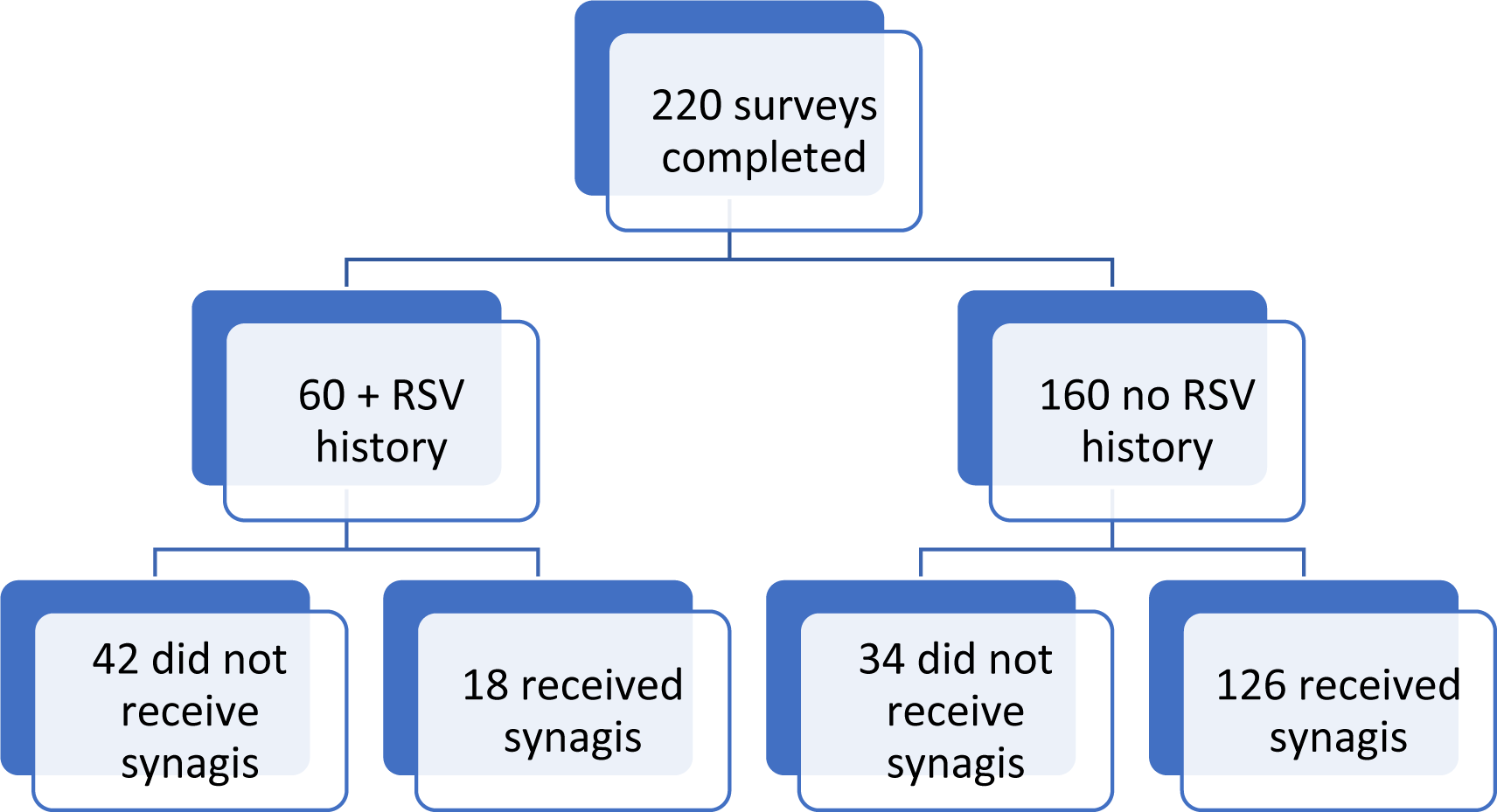
Characteristics of Patients.

Twenty children required CPAP during the infection and 46 children required supplemental oxygen during admission. It was reported that only 8 children who contracted RSV had a history of chronic lung disease prior to the RSV infection (13%) and only 2 had a prior history of asthma (3%), but these individuals required longer hospitalizations with more intensive care. Interestingly, 42% of the patients who had RSV were over age 1 year at the time of infection and 17 of these (68%) were over 24 months of age when they contracted RSV. None of these individuals met the criteria for the Synagis vaccine, and, thus, did not receive it. Fifteen individuals with PWS had multiple RSV infections in childhood, and required recurrent hospitalization. Seventeen patients had chronic lung conditions after the RSV infection. The case lethality was 1.37%.

Respondents stated that 84 of the children with PWS received the Synagis shot at less than 1 year of age (38%) and 41 of those received more than one season of the vaccination. Repeated administration of the vaccine over more than one RSV season was done at the insistence of the parents, not recommendations of the provider. Parents reported that 28% of their babies with PWS were born prematurely and 99% reported that their child had significant hypotonia. Of those individuals who never had RSV, 78.8% received the Synagis vaccine (p<0.001) (Figure 1). Forty-two of the 60 children who had RSV (70%) had not received Synagis (p<0.01) (Figure 1). As expected, RSV infections were most prevalent during the months of December to March.

Parents reported various reasons for their child with PWS not receiving the vaccine, but approximately 30% sited failure of insurance to provide authorization for the treatment or failure of the physician to recommend the treatment. Parents commented how frustrated and anxious they were when they knew their child was at high risk for complications from RSV due to hypotonia, but were unable to get this “potentially life-saving vaccine”.

## CONCLUSIONS

Children with PWS are at high risk for mortality from respiratory infections. This risk is likely due to their hypotonia and inability to adequately cough to clear respiratory secretions [8]. Our survey showed that receiving the Synagis vaccine was associated with a lower incidence of contracting RSV, whereas those who were unable to get the vaccine had a significantly higher risk of contracting the virus. Children with PWS who had RSV had a high rate of hospitalization and need for intensive care. Some of the children who were hospitalized required prolonged hospitalizations or repeat hospitalizations, thus increasing their medical care cost. Of particular note in this syndrome, is that a high percentage of those who contract RSV do so after 24 months of age. All of those who were over 24 months of age required hospitalization due to this infection. Therefore, it is important to consider vaccination for children with PWS of any age who still have significant hypotonia.

This study had several limitations. Response bias may have affected data consistency and no confirmation from medical records was collected. Although the registration form implied that one parent of a child with PWS complete the survey, no digital control for duplicates was established. The study was cross-sectional and not longitudinal, and did not include families who had not consented to participate in research via the support groups.

Despite the limitations, we feel that the information obtained from this survey demonstrates that it would be in the best interest of insurance companies to change their coverage for Synagis for individuals with PWS, and approve coverage for patients of any age who have significant hypotonia with this syndrome.

## REFERENCES

1. Cassidy SB, Schwartz S, Miller JL, Driscoll DJ. Prader-Willi syndrome. Genet Med. 2012 Jan;14(1):10–26.

2. Duis J, van Wattum PJ, Scheimann A, Salehi P, Brokamp E, Fairbrother L, et al. A Multidisciplinary Approach to the Clinical Management of Prader-Willi Syndrome. Mol Genet Genomic Med. 2019 Mar;7(3):e514

3. Butler MG, Manzardo AM, Forster JL. Prader-Willi Syndrome: Clinical Genetics and Diagnostic Aspects With Treatment Approaches. Curr Pediatr Rev. 2016;12(2):136–66.

4. Kimonis VE, Tamura R, Gold JA, Patel N, Surampalli A, Manazir J, Miller JL, Roof E, Dykens E, Butler MG, Driscoll DJ. Early Diagnosis in Prader-Willi Syndrome Reduces Obesity and Associated Co-Morbidities. Genes (Basel). 2019 Nov 6;10(11):898.

5. Mirra V, Ullmann N, Cherchi C, Onofri A, Paglietti MG, Cutrera R. Respiratory Syncytial Virus Prophylaxis and the “Special Population”. Minerva Pediatr. 2018 Dec;70(6):589–599

6. Pacoricona Alfaro DL, Lemoine P, Ehlinger V, et al. Causes of death in Prader-Willi syndrome: lessons from 11 years’ experience of a national reference center. Orphanet J Rare Dis. 2019;14(1):238.

7. Butler MG, Manzardo AM, Heinemann J, Loker C, Loker J. Causes of Death in Prader-Willi Syndrome: Prader-Willi Syndrome Association (USA) 40-year Mortality Survey. Genet Med. 2017 Jun;19(6):635–642.

8. Manzardo AM, Loker J, Heinemann J, Loker C, Butler MG. Survival Trends From the Prader-Willi Syndrome Association (USA) 40-year Mortality Survey. Genet Med. 2018 Jan;20(1):24–30.

9. Chan M, Park JJ, Shi T, Martinón-Torres F, Bont L, Nair H; Respiratory Syncytial Virus Network (ReSViNET). The Burden of Respiratory Syncytial Virus (RSV) Associated Acute Lower Respiratory Infections in Children With Down Syndrome: A Systematic Review and Meta-Analysis. J Glob Health. 2017 Dec;7(2):020413.

